# Genetic Circuits Combined with Machine Learning Provides Fast Responding Living Sensors

**DOI:** 10.1101/2020.10.29.361220

**Authors:** Behide Saltepe, Eray Ulaş Bozkurt, Murat Alp Güngen, A. Ercüment Çiçek, Urartu Özgür Şafak Şeker

## Abstract

Whole cell biosensors (WCBs) have become prominent in many fields from environmental analysis to biomedical diagnostics thanks to advanced genetic circuit design principles. Despite increasing demand on cost effective and easy-to-use assessment methods, a considerable amount of WCBs retains certain drawbacks such as long response time, low precision and accuracy. Furthermore, the output signal level does not correspond to a specific analyte concentration value but shows comparative quantification. Here, we utilized a neural network-based architecture to improve the aforementioned features of WCBs and engineered a gold sensing WCB which has a long response time (18 h). Two Long-Short Term-Memory (LSTM)-based networks were integrated to assess both ON/OFF and concentration dependent states of the sensor output, respectively. We demonstrated that binary (ON/OFF) network was able to distinguish between ON/OFF states as early as 30 min with 78% accuracy and over 98% in 3 h. Furthermore, when analyzed in analog manner, we demonstrated that network can classify the raw fluorescence data into pre-defined analyte concentration groups with high precision (82%) in 3 h. This approach can be applied to a wide range of WCBs and improve rapidness, simplicity and accuracy which are the main challenges in synthetic biology enabled biosensing.

## Introduction

Biosensors are devices composed of biological components (i.e., enzymes^1–5^, tissues, antibodies^6^, nucleic acids^7^, or cells^8–10^) that detect analytes of interest. Developments in recombinant DNA technologies and synthetic biology have increased the design of living biosensors in many fields from medical^11–18^ to environmental applications^18–25^. Besides, they have plenty of advantages over other types of sensors since they are easy-to-use, cost-effective, renewable and very selective^26,27^.

Contrary to well-established analysis methods such as high-performance liquid chromatography (HPLC), or mass spectroscopy which is used to determine amounts of contaminants in samples, whole cell biosensors (WCBs) detect the bioavailable fraction of chemicals. Hence, WCBs are highly sensitive towards tested chemicals be detected at much lower concentrations^28,29^. Furthermore, real-time monitoring of bioavailability of chemicals is possible^30^. Additionally, the exploitation of specific transcription factors (TFs) in circuits make WCBs highly selective allowing the detection of the target compound. Also, combination of different TFs allows construction of multifunctional WCBs to sense and report the presence of multiple molecules simultaneously^29–31^.

Despite successful laboratory results and the aforementioned advantages, only a limited number of WCBs have been adapted to market because of several challenges such as (i) detection of diffusible substances, (ii) slower response compared to conventional protein-based biosensors, (iii) interference of complex environmental samples, and (iv) cell survival^29,30,32^. To overcome these challenges and make the WCBs affordable for usage in daily life, several methods including chassis engineering^30^, lyophilization^33,34^, cellular immobilization^35–40^, continuous culture allowing constant nutrient supply and real-time monitoring^34,41,42^, or advanced TF engineering tools^43^ to decrease response time and increase sensitivity have been employed.

Selectivity, reproducibility, accuracy and sensitivity characteristics are the core parameters that define the functionality of a WCB^44^. While selectivity is ensured by specific transcription factors; reproducibility, accuracy and sensitivity of the results are error-prone and depend on many factors including the experimenter, time, and growth phase of bacteria which can lead to variations in sensor signal^45,46^. Thus, rigorous processing of the biosensor data is of utmost importance for a WCB^47–49^. Moreover, the input–output signal relationships are seldom linear due to natural limitations of the biological processes in cellular biosensing (e.g. cellular volume, concentration of available reactants, etc.). In order to properly interpret the measurements and obtain coherent results, the data has to be processed with techniques that compensate for the non-linearity caused by experimental variations. These additional steps require a quantitative (system-level) understanding of how a biosensor works^50^. Mathematical models of the biosensing system can help better understanding the behavior of sensors. However, relevant parameters of the equations describing the system cannot always be accurately determined as they require equipment that may not always be available or they may permanently disrupt the operation of the biosensor^51,52^. Furthermore, interpretation of output signal differs based on the type of the sensor. Digital sensors interpret the analyte in an ON/OFF fashion and conversion of the signal to field application testing is straightforward. However, analog sensors respond proportionally to analyte concentrations. Interpretation of the output signal in a concentration dependent manner and conversion of the output to analyte concentration is still one of the major drawbacks of WCBs^18^.

Several attempts have been made to make more convenient biosensing platforms for biomedical and environmental applications and to improve the performance of biosensors. For instance, it is anticipated that integration of wireless technology will ease the biomarker detection and provide real-time monitoring. A recently developed micro-bio-electronic device (IMBED) provides a wireless communication of WCBs with ultralow-power microelectronics technology, enabling real-time monitoring of disease biomarkers in the gastrointestinal tract^12^. On the other hand machine learning has been forecasted to play an important role in advancing biosensing^42^ as neural networks in particular have proven useful in an extremely wide spectrum of applications. Yet, there is no study that utilizes machine learning algorithms to advance WCB development.

Artificial neural networks, are a class of non-linear signal processing algorithms (a part of machine learning) that are used to process data that cannot be analytically solved. Using different learning algorithms to analyze the data, the network can be trained to form its own model to fit the data which could then be used to analyze further samples. Neural networks have been used to process biosensor outputs for several applications^53,54^. For instance, Gutes *et al*. used a neural network to determine the type and concentration of phenolic compounds from a polyphenol oxidase amperometric biosensor^53^. Similarly, Trojanowicz *et al.* used them to detect the presence of pesticides from enzymatic biosensors^54^. Deep neural networks employ more hidden layers (hence deep) to learn a hierarchical representation of the features to solve more complicated tasks^55^. One example of such networks is recurrent neural networks (RNN). In the RNN architecture, the output generated from the earlier inputs are fed to the network as an additional input in the next iteration^55^. This property allows the network to remember its state and alter its weights, during training, to better adapt to the data at hand. RNNs are especially useful for applications that require the analysis of temporal or sequential data like sound recognition or genomic analysis^56,57^. The long-short-term-memory (LSTM) network is a more advanced type of RNN that is widely used in deep learning today. LSTMs overcome some of the issues associated with RNNs, and can keep better track of temporal patterns within the data. LSTM networks are also being widely used to analyze various biological functions like DNA– protein binding predictions^58^.

In this paper, we utilized deep neural networks to analyze the output of a biosensor in order to accurately determine target concentrations in much shorter time than possible via manual analysis. First, we engineered a complex genetic circuit to detect gold ions, and characterized the limit of detection (LoD) and response time of the WCB. The output of the sensor reached the maximum fold change (~10-fold) in 18 h. We utilized an LSTM-based network to decrease the detection time of the sensor and to assess the concentration of analyte accurately. First, we predicted the ON/OFF status of the sensor, and achieved high precision in a short time (78% in 30 min, and 93% in 2 h). Next, we trained a second LSTM model and predicted a concentration for the analyte and observed that the model made precise predictions of concentrations in 3 h. Here, we showed that integration of the machine learning in WCBs can be utilized to decrease long response times of sensors and accurately predict applied gold ion concentrations. This study is unique in the field of WCBs and can be further extended to analyze different parameters that might alleviate the labor-intensive work.

## Results

### Construction and characterization of bacterial gold detecting sensor

Whole-cell biosensors hold great potential in many areas, specifically in environmental contaminant analysis and numerous WCBs have been proposed in last two decades^59^. Due to their certain advantages such as cost, rapidness and ease of use, they become a prominent alternative.

In our engineered bacterial WCB design, we constructed a complex and tightly controlled gold detecting circuit combining a semi-specific stress biosensor based on heat shock response (HSR)^60^ and a specific biosensor for gold sensing^61^. In the circuit, a constitutively expressed HSR repressor, HspR, blocks gene expression from its cognitive promoter, P_dnaK-IR3-IR3_, controlling the expression of gold specific transcription factor, GolS, and a site-specific recombinase, Bxb1. Blocking of both elements ensures the elimination of output expression which requires conversion of gold specific promoter, P_golB_, by the site-specific recombinase, and transcription initiation by GolS-gold ion complex (Fig. 1a). The circuit takes action only when gold ions are introduced to the environment causing stress to cells which releases HspR from the HSR promoter and initiates both Bxb1 and GolS expression. First, Bxb1 recognizes certain sequences around the gold specific promoter and converts^62–64^ it towards the output gene. Next, GolS-gold ions complex^19,61^ helps initiate the output expression (Fig. 1b).

**Fig. 1.**
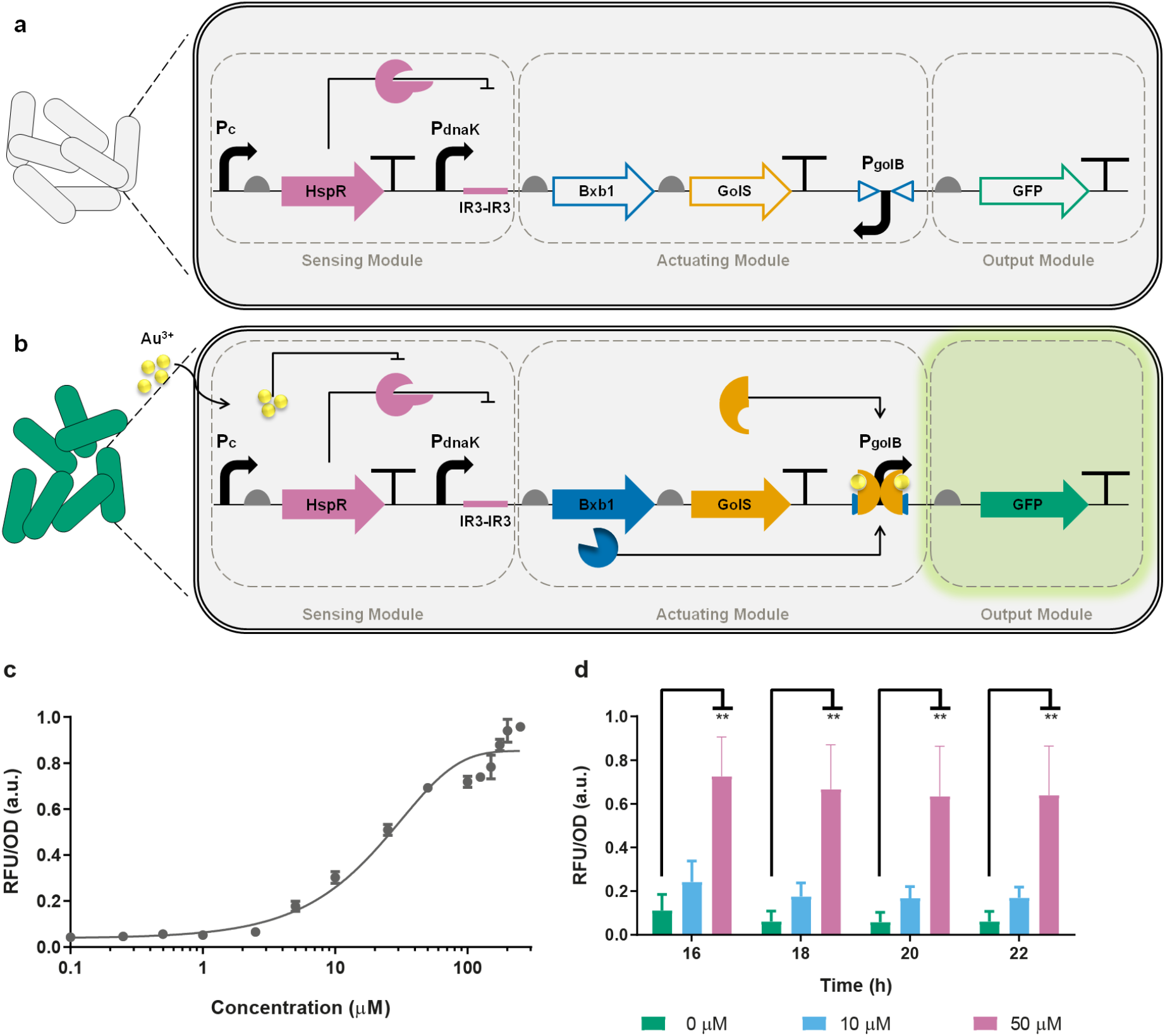
Working principle and characterization of gold sensor. **a,** Without gold, a constitutive promoter (P_c_) expresses HspR repressor. HspR recognizes IR3-IR3 sequences on stress promoter (P_dnaK_) blocking the *bxb1* and *golS* expression. No *gfp* expression is observed from inverted gold specific promoter (P_golB_). **b,** In the presence of gold, HspR releases the stress promoter and Bxb1 and GolS expression is initiated. First, Bxb1 recognizes specific sequences (open blue triangles) and flips gold-specific promoter. Next, GolS-gold complex initiates GFP expression. **c,** Dose response analysis of gold sensor in LB media. 250 μl of cultures in 96-well plates were induced with gold concentrations from 0-to-250 μM and incubated for 18 h at 30°C in a stable incubator. Values are mean ± s.e.m. (n = 6 biologically independent experiments). a.u., arbitrary units. **d,** Time dependent response analysis of gold sensor in MOPS minimal media. 250 μl of cultures in 96-well plates were induced with 0, 10 and 50 μM of gold concentrations and incubated at 30°C in a stable incubator. Values are mean ± s.e.m. (n = 3 biologically independent experiments). a.u., arbitrary units.

To begin with, we optimized the gold detecting WCB response with a dynamic range analysis (Fig. 1c). We induced the sensor with varying gold ion concentrations from 0-to-250 μM for 18 h at 30°C in a stable incubator. We observed that the sensor starts responding to gold ions from 5 μM, which we defined as the LoD of the sensor, the signal increases proportionally with increased gold concentrations, and tends to saturate after 100 μM of gold induction. Therefore, we defined a moderate concentration (i.e. 50 μM) that could be suitable to obtain high response, yet does not disturb cell viability. Next, we examined the specificity of the sensor with 50 μM concentration of varying heavy metal ions (Au^3+^, Cd^2+^, Fe^2+^, Fe^3+^, Co^2+^, Co^3+^, Pb^2+^, As^3+^) and results indicated a significant increase in reporter expression only in gold induced group after 18 h of incubation (Supplementary Figure 2). Lastly, we induced the sensor with low and mid concentrations of gold ions (10 and 50 μM, respectively) to analyze the ideal response time of the sensor (Fig. 1d). We observed that similar responses have been obtained from the sensor within 16 to 22 h of incubation which is quite late for the ideal performance of biosensors. Thus, we introduced LSTM network to decrease the detection times of such biosensors.

**Fig. 2.**
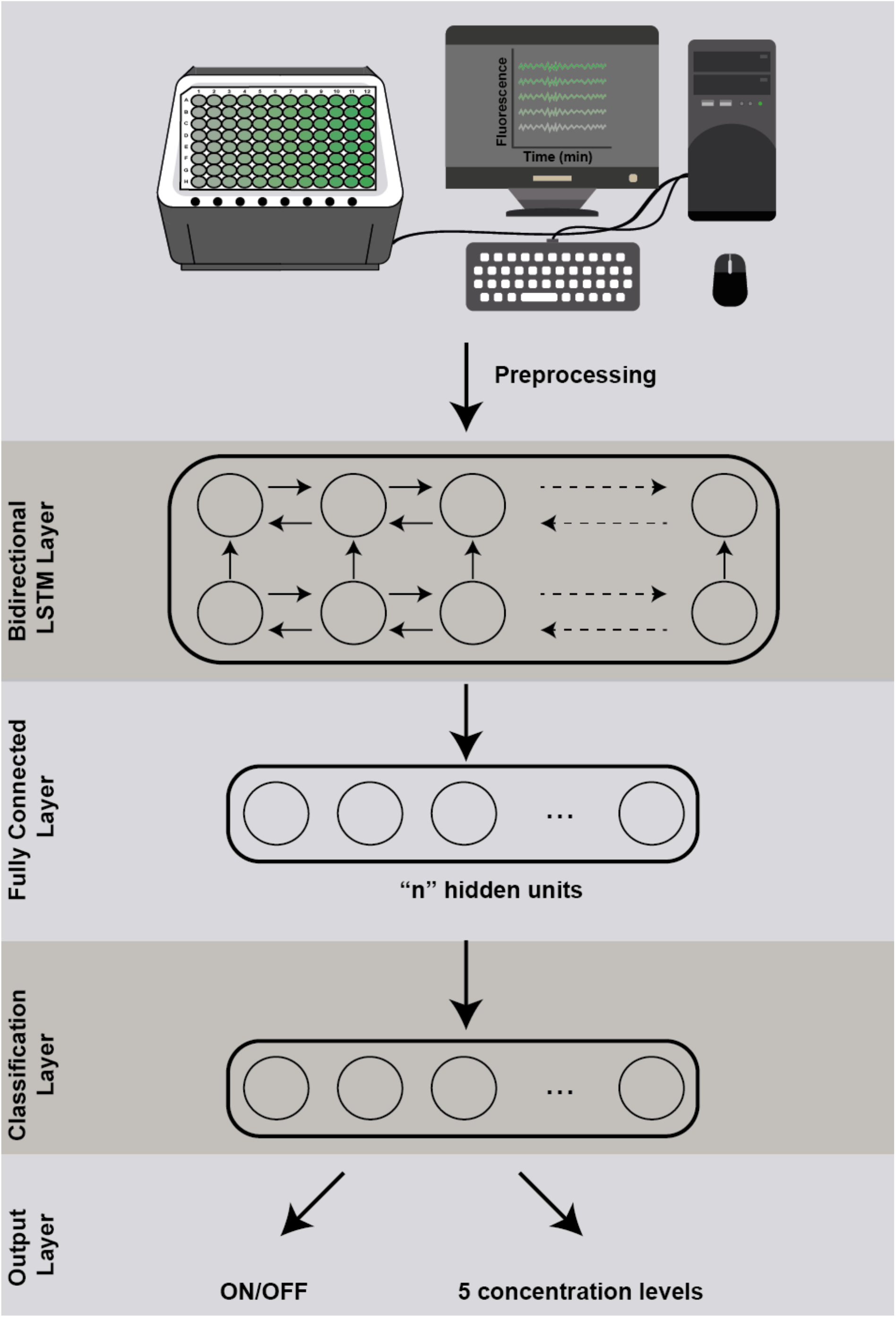
Representation of binary and 5-class neural network architectures. Fluorescence data was measured with 2 min of intervals in a microplate reader for 6h. The raw fluorescence data was pre-processed and input in to a bidirectional LSTM layer (with 90 and 70 hidden units, respectively) which is connected to a fully connected layer with the same number of neurons (2 and 5 hidden units for binary and 5-class classification, respectively) as output classes with softmax activation.

### Workflow of the sensor from wet lab to the LSTM network model

Data processed by the two LSTM networks were obtained from cells induced with gold ions in 96-well plates at 30°C and signal was tracked with 2 min intervals for 6 h. The raw fluorescence data was split into training and test (unseen) data, and processed with LSTM network to shorten the detection time and specify related concentration predictions (Fig. 2). See Methods for details.

### Detection of the presence of gold ions by the LSTM network model

WCBs often suffer from long hours to reach a detectable signal (i.e. 18 h for the highest fold change) which is one of the major disadvantages of WCBs^45,46^. By utilizing an LSTM-based model we aimed to shorten the required time to make an assessment.

In order to optimize and shorten the response time of the sensor, multiple networks were trained and tested for different lengths of the biosensor data (see Methods). Starting from 0 min to 6 h, all data were analyzed with 30 min increments (Fig. 3a). The network was able to identify the ON/OFF states (binary classification) of the sensor in 30 min with high accuracy (78%) and reached the maxima in 3 h (over 98%). The binary classification results of 30 min were represented by confusion matrices for each of the cross validation runs (Fig. 3b). Note that the raw fluorescence signal was not sufficient to show the ON/OFF status of the sensor in 30 min (Fig. 3c), and the earliest discernible appearance of the visual signal (~2-fold signal increase) was observed after 5 h (Supplementary Figure 7.a); however, the LSTM network was able to accurately classify the sensor output. Additionally, binary classification results of 1-to-3 h were represented as confusion matrices in Supplementary Figure 3-5.

**Fig. 3.**
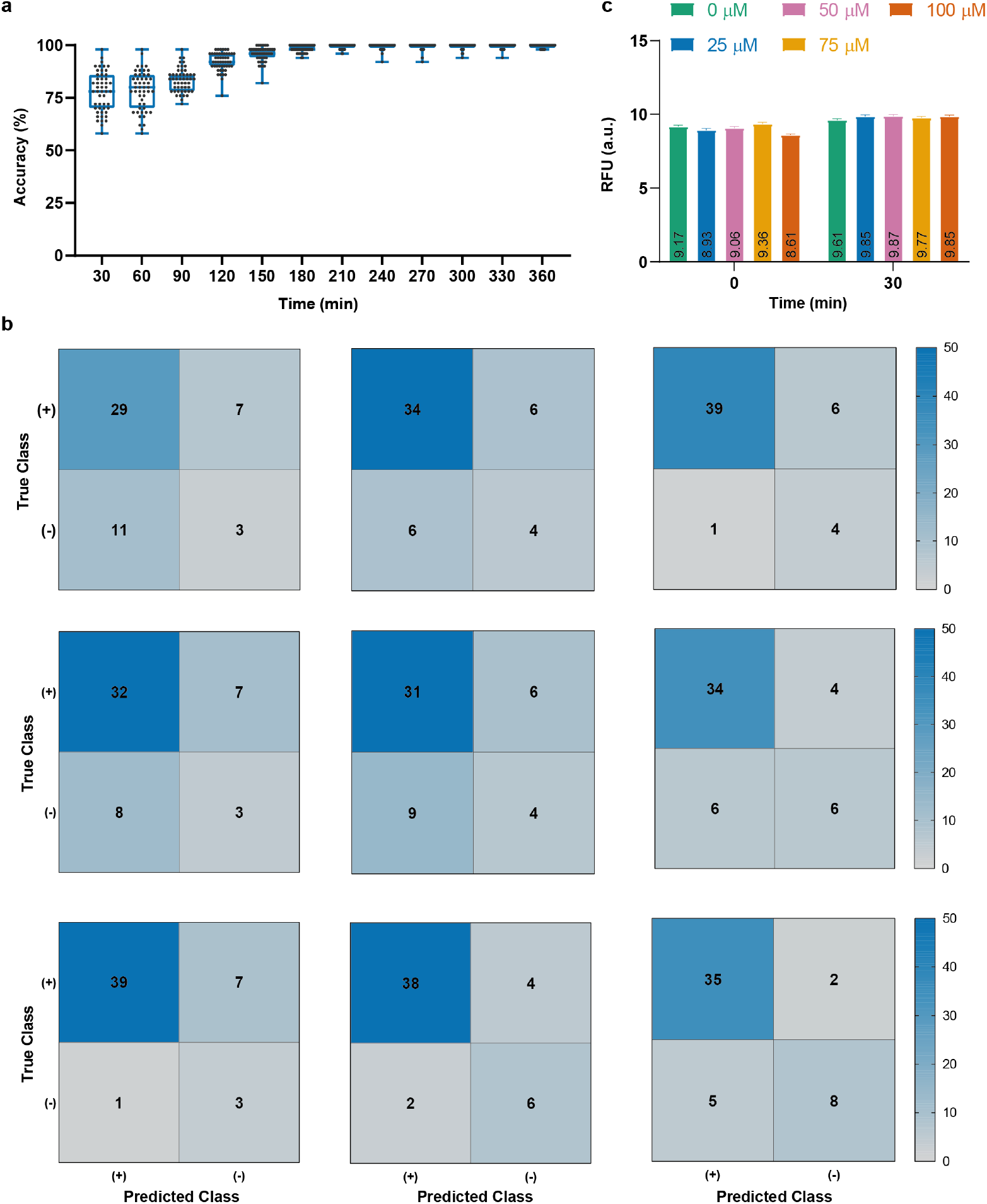
ON/OFF classification accuracy of gold sensor. **a,** The binary (ON/OFF) output of the sensor was achieved with 78% accuracy in the first 30 min, which then increased to over 98% after 3 h, reaching 100% at 6 h. **b,** Confusion matrices of ON/OFF state at 30 min. **c,** Raw fluorescence signals of gold sensor at 0 and 30 min. 250 μl of cultures in 96-well plates were induced with 0, 25, 50, 75 and 100 μM of gold concentrations and incubated at 30°C in a stable incubator. Values are mean ± s.e.m. (87 ≤ n ≤ 96 biologically independent experiments). a.u., arbitrary units.

### Classification of gold concentrations by the LSTM network model

Upon satisfactory results of binary classification, we trained another LSTM model to predict the level of the gold ion concentrations (i.e., discrete gold ion concentration ranges). The results showed that the network could accurately classify the data in 3 h (82%) based on gold concentrations (Fig. 4a). Each cross-validation run was represented by confusion matrices (Fig. 4b). Even though signal ratios showed slight differences between applied concentrations in 3 h (Fig. 4c), the results indicated that the prediction ability of the network is highly satisfactory for both detecting the presence of gold ions (binary classification) and concentration dependent (analog) classification.

**Fig. 4.**
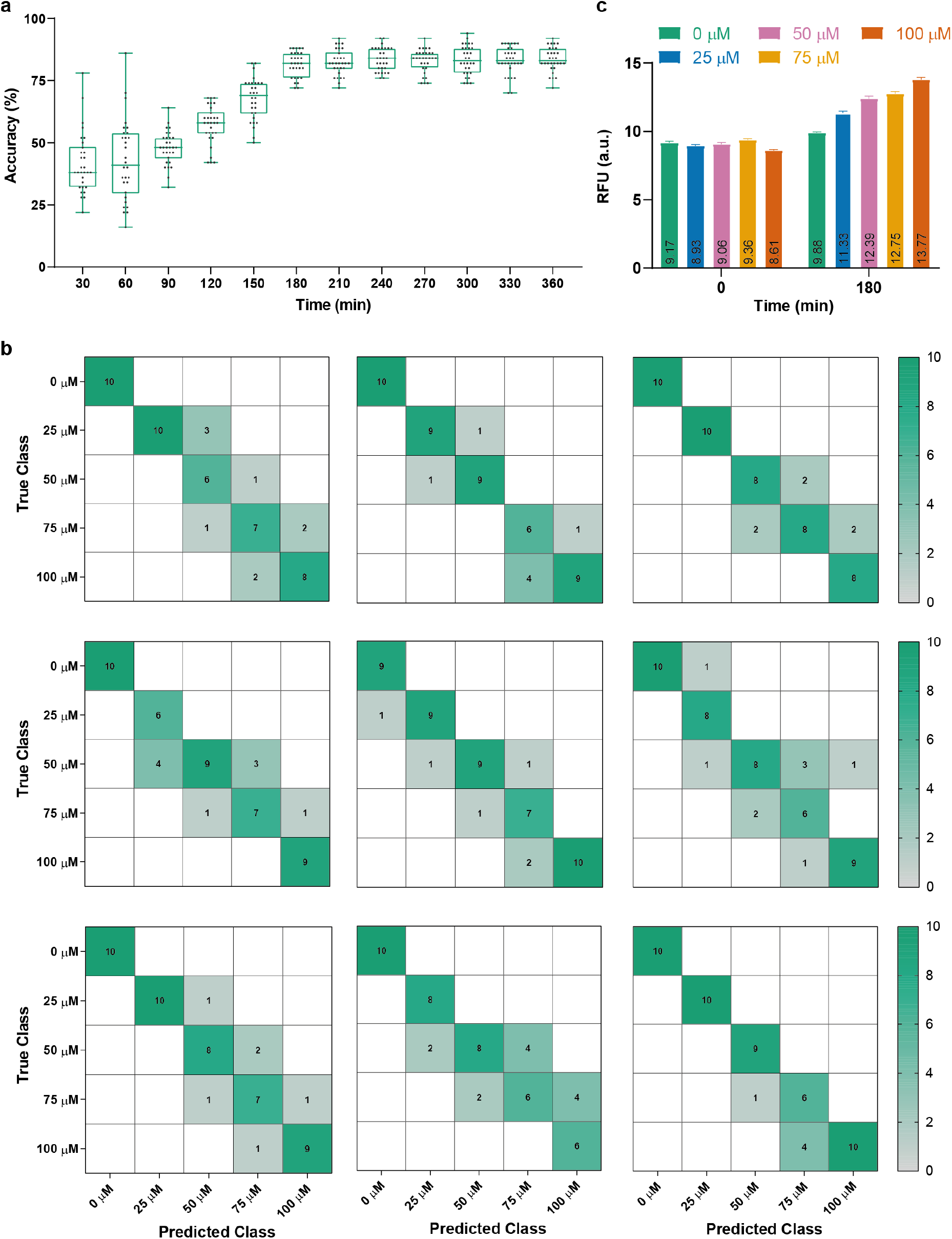
Concentration classification accuracy of gold sensor. **a,** The concentration classification has been reached to 82% in 3 h. **b,** Confusion matrices of concentration classification at 3 h. **c,** Raw fluorescence signals of gold sensor at 0 and 3 h. 250 μl of cultures in 96-well plates were induced with 0, 25, 50, 75 and 100 μM of gold concentrations and incubated at 30°C in a stable incubator. Values are mean ± s.e.m. (87 ≤ n ≤ 96 biologically independent experiments). a.u., arbitrary units.

## Discussion

WCBs are promising tools of biosensing that could be engineered to detect various analytes and could be used in many fields^18^. Yet, there are certain drawbacks to be solved such as long response time. Here, we engineered a gold detecting sensor utilizing gold specific TF, GolS, and a site-specific recombinase, Bxb1. This architecture of the biosensor allows us to monitor both the presence of toxicity as well as the source of the toxicity. The double output capability comes at a cost of longer response time (Figs. 1c and 1d). Therefore, we utilized neural networks to improve the features of our proposed sensor. LSTM is a neural network architecture that is effective in analyzing sequential data^65^. Hence, we chose LSTM as our neural network architecture to learn temporal features from the output of the sensor in relation to the concentration of gold ions.

In our proposed work, we successfully integrated two LSTM-based neural networks to accurately and efficiently predict (i) the presence/absence of gold ions (ON/OFF) and (ii) discrete gold concentration using raw fluorescence signal. The ON/OFF state of the sensors is widely used including pathogen detection and early diagnosis^13,66,67^. Therefore, response time of a WCB is vital for decision-making and the treatment of patients. In this study, we have shown that a machine learning based solution could be integrated to WCBs to decrease the detection time. Although our WCB required 5 h to develop a distinguishable signal (~2-fold) (Supplementary Figure 7), our models were able to shorten the response time to 30 min with 78% accuracy, reaching 93% in 2 h, and over 98% in 3 h (Fig. 3a).

Furthermore, biosensors with analog circuits have been widely used to detect-and-report the presence of an analyte in concentration dependent manner, and play a crucial role in environmental heavy-metal detection^21,43^. Although, WCBs with analog circuits provide quantitative analysis, no study has reported a direct relation between the raw reporter signal and the analyte concentration. Nevertheless, in our study, the LSTM-based architecture was able to classify the data based on the reporter signal behavior utilizing pre-defined concentration classes. Processing of gold-sensing WCB data with another LSTM-based network showed that conversion of raw signal to a discrete concentration value is possible and data can be classified accurately (82%) in 3 h (Fig. 4). Our results showed that the utilization of this model to process WCB data is favorable in terms of (i) decreasing response time, (ii) providing a simple output (rather than a raw fluorescence data), and (iii) allowing concentration classification of a single sample based on the signal.

We envision that this approach can be further optimized to calculate the exact analyte concentration rather than to classify it. To do so, the dynamic range should be explicit and pre-defined concentration groups should be selected from analog region of dynamic range in order to get accurate results. Alternatively, the data from both analog and digital response can be trained with the LSTM model so that a better fit can be obtained for a wider range of concentrations. The last but not the least, this approach can be purposed to decrease the LoD which is one of the main challenges in biosensors. In this study, we defined the LoD of the WCB as 5 μM based on dynamic range analysis (Fig. 1c) which shows only ~2-fold increase in 18 h while we were able to decrease the detection time to 6 h with 75% accuracy (Supplementary Figure 6) with the help of LSTM model.

Engineering the WCB circuits or growth conditions of cells might change the network architecture. For instance, utilization of other reporters (i.e. enzymes) can directly affect the model, since they have different characteristics of signal accumulation and response times. Similarly, the nutrient-rich media could boost the signal accumulation resulting in faster response. Especially, in some cases, introducing additional genetic parts (i.e. recombinases, multiple TFs) to circuits result in a significant delay in response time. Although these tools bring certain advantages such as specificity, they become insufficient tools for biosensing because of the delay. Integration of machine learning based algorithms can benefit such studies.

After creating a specific neural network model for a biosensor, the trained model can be incorporated into portable microcontroller or field programmable gate array (FPGA) based systems. Such platforms can be combined with onboard portable spectrophotometers to provide on-site measurements. This enables obtaining rapid, simple and accurate results in the field with low cost equipment. Given the current situation of the COVID-19 pandemic, the importance of developing such systems, especially to monitor the presence of biomarkers for pathogens has a critical importance for a better healthcare system.

## We thank

We thanks TUBITAK Grant No 114Z653 and 118S398. UOSS and AEC acknowledge the support of TUBA GEBIP and Bilim Akademisi BAGEP awards.

## Author contributions

UOSS conceived the idea, UOSS and AEC designed the study, BS, EUB and MG carried out the experiments. All of the authors wrote the paper.

## Additional information

**Supplementary information** is available for this paper.

**Correspondence and requests for materials** should be addressed to U.O.S.S. and A.E.C

**Publisher’s note:** Springer Nature remains neutral with regard to jurisdictional claims in published maps and institutional affiliations.

## Methods

### Strains, plasmids and bacterial cultivation conditions

*E. coli* DH5α cells were used for both plasmid construction and reporter expression assays. Cells were cultivated in Luria-Bertani (LB) medium (10 g/l tryptone, 5 g/l yeast extract, 5 g/l NaCl) with proper antibiotics (34 μg/ml chloramphenicol stock, 100 μg/ml ampicillin stock). Overnight cultures were prepared from frozen glycerol stocks in 2 ml LB, and cultivated at 37°C with shaking (180 r.p.m.) (INNOVA 44, New Brunswick Scientific). To start experimental cultures, 0.4% of inoculums from overnight cultures were diluted in fresh LB or MOPS minimal media (0.1 M potassium morpholinopropane sulfonate (MOPS), pH 7.4; 0.1 M Tricine, pH 7.4; 0.001 M FeSO_4_; 0.19 M NH_4_Cl; 0.0276 M K_2_SO_4_; 0.002 CaCl_2_; 0.25 M MgCl_2_; 0.5 M NaCl; micronutrients [3×10^−2^ M (NH_4_)_6_Mo_7_O_24_; 4×10^−5^ M H_3_BO_3_; 3×10^−6^ M CoCl_2_; 10^−6^ M CuSO_4_; 8×10^−6^ M MnCl_2_; 10^−6^ M ZnSO_4_]; 0.132 M K_2_HPO_4_; 1 mg/ml thiamine; 0.2% (v/v) glucose) defined by ref.^68^. Each culture was inoculated in 96-well microplates (353916, Corning) to final volume of 250 μl per well and induced with proper inducers. Cells were incubated at 30°C for 18 h in a stable incubator (INCU-Line, VWR), unless otherwise stated. After 18 h of incubation, both cell growth and GFP fluorescence were monitored using microplate reader (SpectraMax M5, Molecular Devices).

Antibiotics and other chemicals used for induction assays (FeSO_4_•7H_2_O, FeCl_3_•6H_2_O, Na_3_AsO_4_, CoCl_2_•6H_2_O, Co(NH_3_)_6_Cl_3_, PbCl_2_, HAuCl_4_, Cd(OOCCH_3_)_2_•2H_2_O) were analytical grade and purchased from Sigma-Aldrich. Each reagent was dissolved in ddH_2_O and filter sterilized using 0.22 μm syringe filters (16532K, Sartorius AG).

### Sensor plasmid construction

Gold sensor plasmids were constructed using standard molecular biology techniques. Primers used in this study were listed in Supplementary Table 1 and purchased from PRZ BioTech. All genetic parts used in this study were summarized in Supplementary Table 2. Plasmid map representations were provided in Supplementary Figure 1. All plasmids were verified by Sanger sequencing (GENEWIZ).

To construct sensing and output modules, inverted PgolB promoter with Bxb1 recognition sites and reporter-terminator (GFP-rrnBT1) pair were amplified by polymerase chain reaction (PCR) using Q5 High-Fidelity DNA Polymerase (M0491, NEB) in thermal cycler (C1000 Touch, Bio-Rad). For backbone, formerly constructed mProD HspR pET22b vector^60^ was linearized with SpeI restriction endonuclease enzyme (R3133, NEB). To construct actuating module, Bxb1 recombinase and GolS were amplified by PCR. For backbone, formerly constructed PdnaK-IR3-IR3 GFP pZa vector^60^ was digested with MluI restriction endonuclease enzyme (R3198, NEB) to exclude *gfp* from the vector. All pieces were run on 1 or 2% (w/v) Agarose gel stained with SYBR Safe DNA gel stain (S33102, Invitrogen). Bands at expected sizes were isolated from the gel (740609.50, MN), and concentrations were quantified with spectrophotometer (NanoDrop 2000, Thermo Fisher Scientific). Both circuits were assembled with Gibson Assembly method^69^. After assembly, entire mixes were transformed into chemically competent *E. coli* DH5α cells.

### Reporter expression assays and data analysis

All assay conditions were described above. Fluorescence for *gfp* expression (485 nm for excitation, 538 nm for emission, 530 nm cut off) and absorbance for optical cell density (OD_600_) were measured via microplate reader. All sensor output was normalized to cell density (*gfp* fluorescence/OD_600_) at specific time point and negative control group (GFP-free cells) was subtracted. Obtained data was normalized in 0-to-1 range: Minimum value was subtracted from each value and divided by the difference between maximum and minimum values (Except Fig. 3c, Fig. 4c, and Supplementary Figure 7).

Continuous GFP expression to feed neural network was measured as following: O/N culture of cells with gold-sensing circuits was diluted 0.4% in fresh MOPS media and placed in a microplate reader. Reporter expression was recorded with 2 min of intervals for 6 h. The experiment temperature was set to 30°C throughout the measurements.

Data was visualized with mean ± standard error mean (s.e.m.) in each graph. At least three biological replicates were used for each analysis. For statistical analysis, one-way analysis of variance (ANOVA) or two-way ANOVA with Dunnett’s multiple comparison tests were used, based on the group of interest. The data was visualized with GraphPad Prism v8 and/or Adobe Illustrator 2015.

### Neural network experimental setup

Two LSTM networks, one with a binary output and the other with five output classes, were implemented using the Deep Learning Toolbox on MatLab (R2019b). In both networks, the raw fluorescence data was preprocessed via the time derivative. The preprocessed data and the raw data were then fed into the network as input. The datasets of both networks used the same set of measurements: 0 μM, 25 μM, 50 μM, 75 μM, and 100 μM, with 87 ≤ n ≤ 96 sample sizes.

Time differentiation was implemented on MatLab according to the following equation. *F* is the vector containing the raw fluorescence measurements and n is the number of elements within the vector.

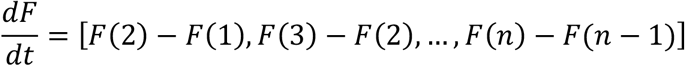

Prior to training, for both networks, the datasets were first split randomly distributed subsets. The same subsets were then used to train and evaluate the results of all iterations of their respective networks. 70% of the data was used for training, 20% for validation and 10% for testing.

### Neural network architecture

For the binary network, the dataset was split such that the 0 μM inputs were labeled as *OFF* (“0” in the network) and the others (25 μM, 50 μM, 75 μM, and 100 μM inputs) were all labeled as *ON* (“1” in the network). The architecture of the network consisted of the following layers: a sequential input layer (of size 2; for the raw data and its time-differentiated counterpart), a bidirectional LSTM layer (using the default *tanh* activation functions in the toolbox), a fully connected layer with two output neurons (each neuron corresponding to one of the outputs), a *softmax* layer (used to implement the activation functions of the fully connected layer), and a classification layer.

In the 5-class (concentration range classifying) network, there were five output classes in total, corresponding to each concentration input (0 μM, 25 μM, 50 μM, 75 μM, and 100 μM). The outputs were in a binary vector format where the positive class was represented by a “1” and the others by “0”. The architecture of the network consisted of the following layers: a sequential input layer (of size 2; for the raw data and its time-differentiated counterpart), a bidirectional LSTM layer (using the default *tanh* activation functions in the toolbox), a fully connected layer with five output neurons (each neuron corresponding to one of the outputs), a *softmax* layer (used to implement the activation functions of the fully connected layer), and a classification layer.

### Neural network optimization

Optimization was performed by altering the following network variables in the presented order; the number of hidden units in the bidirectional LSTM layer, the number of neurons in the number of fully connected layers, the number of neurons in each fully connected layer, and the output functions of each of these layers. Starting from ten, with increments of ten, up to 120, the number of neurons in the bidirectional LSTM layer was altered in both networks. This was followed by experimenting with different numbers of fully connected layers (up to three), number of neurons in each fully connected layer (starting with the same number of neurons as the number of output classes and going up to twenty), and different activation functions for the neurons in each layer (*softmaxLayer, reluLayer*, and *leakyreluLayer*).

Prior to training, the weights were initialized using the MatLab Deep Learning Toolbox’s default *glorot* (Xavier) weight initialization function. Training was performed using the ADAM optimizer with its default learning rate of 0.001. Batch was performed using mini-batches with 200 samples and 700 epochs (leading to a maximum of 1400 iterations). At the start of every epoch, the elements within the batch were shuffled.

### Final Neural network architecture

The binary classifying network has a sequential input layer (of size 2; for the raw data and its time-differentiated counterpart), a bidirectional LSTM layer (using the default *tanh* activation functions in the toolbox) with 90 hidden units, a fully connected layer with two output neurons (each neuron corresponding to one of the outputs), a *softmax* layer (used to implement the activation functions of the two neurons in the fully connected layer), and a classification layer which outputs “1” or “0” corresponding to the “ON” or “OFF” states of the network respectively.

The concentration classifying network has a sequential input layer (of size 2; for the raw data and its time-differentiated counterpart), a bidirectional LSTM layer (using the default *tanh* activation functions in the toolbox) with 70 hidden units, a fully connected layer with five output neurons (each neuron corresponding to one of the outputs), a *softmax* layer (used to implement the activation functions of the five neurons in the fully connected layer), and a classification layer which outputs one of the five possible concentration values (0 μM, 25 μM, 50 μM, 75 μM, and 100 μM).

### Neural network testing

After each network was optimized with respect to the validation subset, the validation subset was combined with the training set and the network was trained from scratch with the expanded training set (consisting of 90% of the data). The previously unseen testing set (containing the remaining 10%) was then used to evaluate the performance of the network.

The same networks were then used for their respective temporal accuracy analysis. In these tests, the length of each element in the dataset was cropped to its respective time-length. For instance, only the measurements for the first 30 min were used initially, followed by longer sequences corresponding to 60, 90, 120 min, etc. In both cases, the accuracy of the networks was determined by comparing the network outputs of the testing set with their true values. In order to achieve coherent results, leave-one-out cross-validation was used. Ten different networks, were trained and tested with different members of the data in the training and testing sets. The overall percentage accuracy was determined as follows:

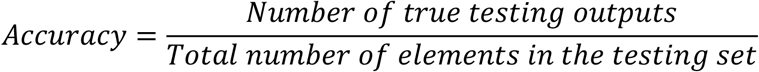

The test result for the temporal accuracy has been plotted in figures 3a and 4a for the binary and 5-class cases respectively. As can be seen from the figures, the accuracy of the results increases alongside the duration of the data. Improving the performance of the system may be possible by measuring the fluorescence of the samples with a higher sampling rate.

## References

1 Akyilmaz, E., Yorganci, E. & Asav, E. Do copper ions activate tyrosinase enzyme? A biosensor model for the solution. Bioelectrochemistry 78, 155–160, doi:10.1016/j.bioelechem.2009.09.007 (2010).

2 Wang, J. Electrochemical glucose biosensors. Chem Rev 108, 814–825, doi:10.1021/cr068123a (2008).

3 Ghasemi-Varnamkhasti, M. et al. Monitoring the aging of beers using a bioelectronic tongue. Food Control 25, 216–224, doi:10.1016/j.foodcont.2011.10.020 (2012).

4 Mishra, R. K., Dominguez, R. B., Bhand, S., Munoz, R. & Marty, J. L. A novel automated flow-based biosensor for the determination of organophosphate pesticides in milk. Biosens Bioelectron 32, 56–61, doi:10.1016/j.bios.2011.11.028 (2012).

5 Chambers, C. E., Visser, M. B., Schwab, U. & Sokol, P. A. Identification of N-acylhomoserine lactones in mucopurulent respiratory secretions from cystic fibrosis patients. Fems Microbiol Lett 244, 297–304, doi:10.1016/j.femsle.2005.01.055 (2005).

6 Conroy, P. J., Hearty, S., Leonard, P. & O’Kennedy, R. J. Antibody production, design and use for biosensor-based applications. Semin Cell Dev Biol 20, 10–26, doi:10.1016/j.semcdb.2009.01.010 (2009).

7 Wang, J. DNA biosensors based on peptide nucleic acid (PNA) recognition layers. A review. Biosens Bioelectron 13, 757–762, doi:Doi 10.1016/S0956-5663(98)00039-6 (1998).

8 Arora, P., Sindhu, A., Dilbaghi, N. & Chaudhury, A. Biosensors as innovative tools for the detection of food borne pathogens. Biosens Bioelectron 28, 1–12, doi:10.1016/j.bios.2011.06.002 (2011).

9 Ercole, C., Del Gallo, M., Mosiello, L., Baccella, S. & Lepidi, A. Escherichia coli detection in vegetable food by a potentiometric biosensor. Sensor Actuat B-Chem 91, 163–168, doi:10.1016/S0925-4005(03)00083-2 (2003).

10 Torun, O., Boyaci, I. H., Temur, E. & Tamer, U. Comparison of sensing strategies in SPR biosensor for rapid and sensitive enumeration of bacteria. Biosens Bioelectron 37, 53–60, doi:10.1016/j.bios.2012.04.034 (2012).

11 Riglar, D. T. et al. Engineered bacteria can function in the mammalian gut long-term as live diagnostics of inflammation. Nat Biotechnol 35, 653–+, doi:10.1038/nbt.3879 (2017).

12 Mimee, M. et al. An ingestible bacterial-electronic system to monitor gastrointestinal health. Science 360, 915–918, doi:10.1126/science.aas9315 (2018).

13 Courbet, A., Endy, D., Renard, E., Molina, F. & Bonnet, J. Detection of pathological biomarkers in human clinical samples via amplifying genetic switches and logic gates. Sci Transl Med 7, doi:ARTN 289ra83 10.1126/scitranslmed.aaa3601 (2015).

14 Watstein, D. M. & Styczynski, M. P. Development of a Pigment-Based Whole-Cell Zinc Biosensor for Human Serum. Acs Synth Biol 7, 267–275, doi:10.1021/acssynbio.7b00292 (2018).

15 Duan, F. P. & March, J. C. Engineered bacterial communication prevents Vibrio cholerae virulence in an infant mouse model. P Natl Acad Sci USA 107, 11260–11264, doi:10.1073/pnas.1001294107 (2010).

16 Hwang, I. Y. et al. Reprogramming Microbes to Be Pathogen-Seeking Killers. Acs Synth Biol 3, 228–237, doi:10.1021/sb400077j (2014).

17 Ho, C. L. et al. Engineered commensal microbes for diet-mediated colorectal-cancer chemoprevention. Nat Biomed Eng 2, 27–37, doi:10.1038/s41551-017-0181-y (2018).

18 Saltepe, B., Kehribar, E. S., Yirmibesoglu, S. S. S. & Seker, U. O. S. Cellular Biosensors with Engineered Genetic Circuits. Acs Sensors 3, 13–26, doi:10.1021/acssensors.7b00728 (2018).

19 Cerminati, S., Soncini, F. C. & Checa, S. K. Selective Detection of Gold Using Genetically Engineered Bacterial Reporters. Biotechnol Bioeng 108, 2553–2560, doi:10.1002/bit.23213 (2011).

20 Wang, B. J., Barahona, M. & Buck, M. A modular cell-based biosensor using engineered genetic logic circuits to detect and integrate multiple environmental signals. Biosens Bioelectron 40, 368–376, doi:10.1016/j.bios.2012.08.011 (2013).

21 Stocker, J. et al. Development of a set of simple bacterial biosensors for quantitative and rapid measurements of arsenite and arsenate in potable water. Environ Sci Technol 37, 4743–4750, doi:10.1021/es034258b (2003).

22 de Mora, K. et al. A pH-based biosensor for detection of arsenic in drinking water. Anal Bioanal Chem 400, 1031–1039, doi:10.1007/s00216-011-4815-8 (2011).

23 Cao, Y. X. L. et al. Programmable assembly of pressure sensors using pattern-forming bacteria. Nat Biotechnol 35, 1087-+, doi:10.1038/nbt.3978 (2017).

24 Zhang, F. Z., Carothers, J. M. & Keasling, J. D. Design of a dynamic sensor-regulator system for production of chemicals and fuels derived from fatty acids. Nat Biotechnol 30, 354–U166, doi:10.1038/nbt.2149 (2012).

25 Belkin, S. et al. Remote detection of buried landmines using a bacterial sensor. Nat Biotechnol 35, 308–310, doi:10.1038/nbt.3791 (2017).

26 Kim, H. J., Jeong, H. & Lee, S. J. Synthetic biology for microbial heavy metal biosensors. Anal Bioanal Chem 410, 1191–1203, doi:10.1007/s00216-017-0751-6 (2018).

27 van der Meer, J. R. & Belkin, S. Where microbiology meets microengineering: design and applications of reporter bacteria. Nat Rev Microbiol 8, 511–522, doi:10.1038/nrmicro2392 (2010).

28 Wei, H., Ze-Ling, S., Le-Le, C., Wen-hui, Z. & Chuan-Chao, D. Specific detection of bioavailable phenanthrene and mercury by bacterium reporters in the red soil. Int J Environ Sci Te 11, 685–694, doi:10.1007/s13762-013-0216-1 (2014).

29 Gui, Q. Y., Lawson, T., Shan, S. Y., Yan, L. & Liu, Y. The Application of Whole Cell-Based Biosensors for Use in Environmental Analysis and in Medical Diagnostics. Sensors-Basel 17, doi:Artn 1623 10.3390/S17071623 (2017).

30 Bereza-Malcolm, L. T., Mann, G. & Franks, A. E. Environmental Sensing of Heavy Metals Through Whole Cell Microbial Biosensors: A Synthetic Biology Approach. Acs Synth Biol 4, 535–546, doi:10.1021/sb500286r (2015).

31 Hou, Q. H. et al. Detection of bioavailable cadmium, lead, and arsenic in polluted soil by tailored multiple Escherichia coli whole-cell sensor set. Anal Bioanal Chem 407, 6865–6871, doi:10.1007/s00216-015-8830-z (2015).

32 Raut, N., O’Connor, G., Pasini, P. & Daunert, S. Engineered cells as biosensing systems in biomedical analysis. Anal Bioanal Chem 402, 3147–3159, doi:10.1007/s00216-012-5756-6 (2012).

33 Ulitzur, S., Lahav, T. & Ulitzur, N. A novel and sensitive test for rapid determination of water toxicity. Environ Toxicol 17, 291–296, doi:10.1002/tox.10060 (2002).

34 Bjerketorp, J., Hakansson, S., Belkin, S. & Jansson, J. K. Advances in preservation methods: keeping biosensor microorganisms alive and active. Curr Opin Biotech 17, 43–49, doi:10.1016/j.copbio.2005.12.005 (2006).

35 Park, K. S. et al. Immobilization as a technical possibility for long-term storage of bacterial biosensors. Radiat Environ Bioph 44, 69–71, doi:10.1007/s00411-005-0271-1 (2005).

36 Polyak, B., Geresh, S. & Marks, R. S. Synthesis and characterization of a biotin-alginate conjugate and its application in a biosensor construction. Biomacromolecules 5, 389–396, doi:10.1021/bm034454a (2004).

37 Philp, J. C. et al. Whole cell immobilised biosensors for toxicity assessment of a wastewater treatment plant treating phenolics-containing waste. Anal Chim Acta 487, 61–74, doi:10.1016/S0003-2670(03)00358-1 (2003).

38 Axelrod, T., Eltzov, E. & Marks, R. S. Bioluminescent bioreporter pad biosensor for monitoring water toxicity. Talanta 149, 290–297, doi:10.1016/j.talanta.2015.11.067 (2016).

39 Buffi, N. et al. Development of a microfluidics biosensor for agarose-bead immobilized Escherichia coli bioreporter cells for arsenite detection in aqueous samples. Lab Chip 11, 2369–2377, doi:10.1039/c1lc20274j (2011).

40 Guo, K. H. et al. Determination of Gold Ions in Human Urine Using Genetically Engineered Microorganisms on a Paper Device. Acs Sensors 3, 744–748, doi:10.1021/acssensors.7b00931 (2018).

41 Pooley, D. T. et al. Continuous culture of photobacterium. Biosens Bioelectron 19, 1457–1463, doi:10.1016/j.bios.2003.09.003 (2004).

42 Lee, J. H. & Gu, M. B. An integrated mini biosensor system for continuous water toxicity monitoring. Biosens Bioelectron 20, 1744–1749, doi:10.1016/j.bios.2004.06.036 (2005).

43 Wan, X. Y. et al. Cascaded amplifying circuits enable ultrasensitive cellular sensors for toxic metals. Nat Chem Biol 15, 540–+, doi:10.1038/s41589-019-0244-3 (2019).

44 Mehrotra, P. Biosensors and their applications - A review. J Oral Biol Craniofac Res 6, 153–159, doi:10.1016/j.jobcr.2015.12.002S2212-4268(15)00132-3 [pii] (2016).

45 Yagi, K. Applications of whole-cell bacterial sensors in biotechnology and environmental science. Appl Microbiol Biot 73, 1251–1258, doi:10.1007/s00253-006-0718-6 (2007).

46 D’Souza, S. F. Microbial biosensors. Biosens Bioelectron 16, 337–353, doi:Doi 10.1016/S0956-5663(01)00125-7 (2001).

47 Wang, X., Errede, B. & Elston, T. C. Mathematical analysis and quantification of fluorescent proteins as transcriptional reporters. Biophys J 94, 2017–2026, doi:10.1529/biophysj.107.122200 (2008).

48 Hebisch, E., Knebel, J., Landsberg, J., Frey, E. & Leisner, M. High Variation of Fluorescence Protein Maturation Times in Closely Related Escherichia coli Strains. Plos One 8, doi:ARTN e75991 10.1371/journal.pone.0075991 (2013).

49 Costa, K. D., Kleinstein, S. H. & Hershberg, U. Biomedical Model Fitting and Error Analysis. Sci Signal 4, doi:ARTN tr9 10.1126/scisignal.2001983 (2011).

50 Rabner, A. et al. Mathematical Modeling of a Bioluminescent E. Coli Based Biosensor. Nonlinear Anal-Model 14, 505–529, doi:Doi 10.15388/Na.2009.14.4.14471 (2009).

51 Ashyraliyev, M., Fomekong-Nanfack, Y., Kaandorp, J. A. & Blom, J. G. Systems biology: parameter estimation for biochemical models. Febs J 276, 886–902, doi:10.1111/j.1742-4658.2008.06844.x (2009).

52 Neves, S. R. Obtaining and Estimating Kinetic Parameters from the Literature. Sci Signal 4, doi:ARTN tr8 10.1126/scisignal.2001988 (2011).

53 Gutes, A., Cespedes, F., Alegret, S. & del Valle, M. Determination of phenolic compounds by a polyphenol oxidase amperometric biosensor and artificial neural network analysis. Biosens Bioelectron 20, 1668–1673, doi:10.1016/j.bios.2004.07.026 (2005).

54 Trojanowicz, M. Determination of pesticides using electrochemical enzymatic biosensors. Electroanal 14, 1311–1328, doi:Doi 10.1002/1521-4109(200211)14:19/20<1311::Aid-Elan1311>3.0.Co;2-7 (2002).

55 Goodfellow, I., Bengio, Y. & Courville, A. Deep learning.

56 Graves, A., Mohamed, A. R. & Hinton, G. Speech Recognition with Deep Recurrent Neural Networks. Int Conf Acoust Spee, 6645–6649 (2013).

57 Shen, Z., Bao, W. Z. & Huang, D. S. Recurrent Neural Network for Predicting Transcription Factor Binding Sites. Sci Rep-Uk 8, doi:Artn 15270 10.1038/S41598-018-33321-1 (2018).

58 Zhang, Y. Q., Qiao, S. J., Ji, S. J. & Li, Y. Z. DeepSite: bidirectional LSTM and CNN models for predicting DNA-protein binding. Int J Mach Learn Cyb 11, 841–851, doi:10.1007/s13042-019-00990-x (2020).

59 Harms, H., Wells, M. C. & van der Meer, J. R. Whole-cell living biosensors - are they ready for environmental application? Appl Microbiol Biot 70, 273–280, doi:10.1007/s00253-006-0319-4 (2006).

60 Saltepe, B., Bozkurt, E. U., Haciosmanoglu, N. & Seker, U. O. S. Genetic Circuits To Detect Nanomaterial Triggered Toxicity through Engineered Heat Shock Response Mechanism. Acs Synth Biol 8, 2404–2417, doi:10.1021/acssynbio.9b00291 (2019).

61 Checa, S. K. et al. Bacterial sensing of and resistance to gold salts. Mol Microbiol 63, 1307–1318, doi:10.1111/j.1365-2958.2007.05590.x (2007).

62 Ghosh, P., Pannunzio, N. R. & Hatfull, G. F. Synapsis in phage Bxb1 integration: Selection mechanism for the correct pair of recombination sites. J Mol Biol 349, 331–348, doi:10.1016/j.jmb.2005.03.043 (2005).

63 Roquet, N., Soleimany, A. P., Ferris, A. C., Aaronson, S. & Lu, T. K. Synthetic recombinase-based state machines in living cells. Science 353, 363–+, doi:10.1126/science.aad8559 (2016).

64 Siuti, P., Yazbek, J. & Lu, T. K. Synthetic circuits integrating logic and memory in living cells. Nat Biotechnol 31, 448–+, doi:10.1038/nbt.2510 (2013).

65 Hochreiter, S. & Schmidhuber, J. Long short-term memory. Neural Comput 9, 1735–1780, doi:10.1162/neco.1997.9.8.1735 (1997).

66 Saeidi, N. et al. Engineering microbes to sense and eradicate Pseudomonas aeruginosa, a human pathogen. Mol Syst Biol 7, doi:Artn 521 10.1038/Msb.2011.55 (2011).

67 Danino, T. et al. Programmable probiotics for detection of cancer in urine. Sci Transl Med 7, doi:ARTN 289ra84 10.1126/scitranslmed.aaa3519 (2015).

68 Neidhardt, F. C., Bloch, P. L. & Smith, D. F. Culture medium for enterobacteria. J Bacteriol 119, 736–747 (1974).

69 Gibson, D. G. et al. Enzymatic assembly of DNA molecules up to several hundred kilobases. Nat Methods 6, 343–U341, doi:10.1038/Nmeth.1318 (2009).

## References Used only in Methods section

61 Neidhardt, F. C., Bloch, P. L. & Smith, D. F. Culture medium for enterobacteria. J Bacteriol 119, 736–747 (1974).

62 Gibson, D. G. et al. Enzymatic assembly of DNA molecules up to several hundred kilobases. Nat Methods 6, 343–U341, doi:10.1038/Nmeth.1318 (2009).

